# Predicting the impact of rare variants on RNA splicing in CAGI6

**DOI:** 10.1101/2023.06.20.545093

**Authors:** Jenny Lord, Carolina Jaramillo Oquendo, Htoo A. Wai, Andrew G.L Douglas, David J. Bunyan, Yaqiong Wang, Zhiqiang Hu, Zishuo Zeng, Daniel Danis, Panagiotis Katsonis, Amanda Williams, Olivier Lichtarge, Yuchen Chang, Richard D. Bagnall, Stephen M. Mount, Brynja Matthiasardottir, Chiaofeng Lin, Thomas van Overeem Hansen, Raphael Leman, Alexandra Martins, Claude Houdayer, Sophie Krieger, Constantina Bakolitsa, Yisu Peng, Akash Kamandula, Predrag Radivojac, Diana Baralle

## Abstract

**Background:** Variants which disrupt splicing are a frequent cause of rare disease that have been under-ascertained clinically. Accurate and efficient methods to predict a variant’s impact on splicing are needed to interpret the growing number of variants of unknown significance (VUS) identified by exome and genome sequencing. Here we present the results of the CAGI6 Splicing VUS challenge, which invited predictions of the splicing impact of 56 variants ascertained clinically and functionally validated to determine splicing impact.

**Results:** The performance of 12 prediction methods, along with SpliceAI and CADD, was compared on the 56 functionally validated variants. The maximum overall accuracy achieved was 82% from two different approaches, one weighting SpliceAI scores by minor allele frequency, and one applying the recently published Splicing Prediction Pipeline (SPiP). SPiP performed optimally in terms of sensitivity, while an ensemble method combining multiple prediction tools and information from databases exceeded all others for specificity.

**Conclusions:** Several challenge methods equalled or exceeded the performance of SpliceAI, with ultimate choice of prediction method likely to depend on experimental or clinical aims. One quarter of the variants were incorrectly predicted by at least 50% of the methods, highlighting the need for further improvements to splicing prediction methods for successful clinical application.

## Introduction

The diagnosis of rare disorders has been revolutionised in recent years thanks to the availability and widespread adoption of next generation sequencing technologies capable of detecting disease-causing variants. With the ever-decreasing prices of whole-exome sequencing (WES) and whole-genome sequencing (WGS) comes an increased use of these approaches, leading to the detection of more genetic variants than ever before. This brings with it a major challenge in understanding what these variants do, since our ability to detect them has far outstripped our ability to meaningfully interpret their effects, particularly outside of protein coding regions. As a result, even with WGS, around half of patients with rare disorders do not get a diagnosis (Turro et al. 2020; Stranneheim et al. 2021).

While estimates vary widely (Lord and Baralle 2021), it is thought somewhere between 15-60% of disease causing variants affect splicing (Krawczak et al. 1992; López-Bigas et al. 2005). Generally speaking, in diagnostic and research variant prioritisation pipelines, variants which fall within the 2bp canonical splice acceptor or donor sites will be classed as splice-affecting, while variants outside of those small regions are often not assessed for splicing impact. It is common for intronic and synonymous variants to be filtered out, while missense variants are generally assessed for their impact on protein structure and function without consideration for the role they may play in splicing. All of these variant types, however, can and do impact splicing and cause disease. This approach has led to an under-ascertainment of splice-affecting variants clinically (Lord et al. 2019). What is needed, particularly with the increasing use of WGS over WES enabling the detection of far more intronic variants than before, is a way to effectively triage which variants are splice-affecting and which are not.

Currently, under ACMG/AMP guidelines (Richards et al. 2015), *in silico* splicing prediction approaches may be used as supporting evidence for genetic diagnosis if multiple independent tools predict an impact on splicing. Experimental validation of splicing effects using RT-PCR, mini-genes or RNAseq is often required to confidently establish a variant’s impact on splicing, but such approaches are time-consuming and expensive to perform at scale. Recent years have seen an explosion of innovative new approaches to splicing prediction, with many new tools being generated, often utilising machine learning. If a high degree of accuracy and reliability can be obtained from *in silico* approaches, we may be able to move away from requiring experimental confirmations, or at the least, have an efficient method to triage variants most in need of validation. This would require highly accurate algorithms and extensive testing in the clinical setting to give sufficient confidence in these optimal approaches.

The Splicing Variants of Unknown Significance (VUS) challenge in the 6^th^ Critical Assessment of Genome Interpretation (CAGI6) sought to assess splicing prediction accuracy on a set of clinically ascertained, functionally validated variants. This enabled performance comparison of many cutting-edge splicing prediction approaches and gave insights into the types of variants not currently well captured by these methods.

## Methods

### Variant selection and validation

As previously described in Wai et al. 2020 (Wai et al. 2020), a total of 64 variants were ascertained through Wessex Regional Genetics Laboratory in Salisbury (52 variants) or the Splicing and Disease research study (12 variants) at the University of Southampton, ethically approved by the Health Research Authority (IRAS Project ID 49685, REC 11/SC/0269) and by the University of Southampton (ERGO ID 23056). Informed consent was provided for all patients for splicing studies to be conducted. All variants had been, or were undergoing RT-PCR analysis to determine their impact on splicing using RNA from whole blood collected in PAXgene tubes, again as previously described (Wai et al. 2020).

Eight variants were excluded from the final analysis (unable to establish splicing impact before analysis period (n=3), incorrect gene/variant annotations given in the dataset distributed (n=3), variant found to impact gene expression rather than splicing (n=2)), giving a total of 56 variants in the final assessment set (**Supplementary Table 1**), which span a wide range of rare disease and cancer predisposition associations, none of which had had their impact on splicing published previously.

### The Splicing VUS challenge

Variants were distributed as a tab delimited text file including the following information: HGNC identifier, chromosome, position, reference allele, alternative allele, gene and strand. Entrants also had access to 256 previously published variants (Wai et al. 2020) obtained and validated by the same approach to aid in method development/testing.

Challenge participants submitted their entries in the form of tab delimited text files including the variant information, a binary prediction of whether a variant affected splicing or not (1=yes, 0=no), along with a score for the probability of the variant affecting splicing and the level of confidence in the prediction given. All assessments were based on the binary splice-affecting prediction alone.

### Challenge assessment

The performance of each prediction model was assessed by calculating and comparing a series of metrics: overall accuracy, area under the receiver operating characteristic curve (AUC), sensitivity, specificity, positive predictive value (PPV) and negative predictive value (NPV). AUC and confidence intervals (2000 stratified bootstrap replicates) were calculated using the pROC package (Robin et al. 2011) in R v3.5.1 (R Core Team 2018), and plots made with ggplot2 (Wickham 2009). Performance of each method was compared across binned splicing locations – Near Acceptor (acceptor +/- 10bp), Near Donor (donor +/- 10bp), Exonic Distant (exonic, 11bp or more from either splice site), Intronic Distant (intronic, 11bp or more from either splice site. For grouped analyses, exonic distant and intronic distant variants were grouped together due to low numbers). These scores were based on the concordance of the binary classification of the variants provided by each team/model (1=splice-affecting and 0=not splice-affecting) with the experimental validation of the splicing impact.

SpliceAI (Jaganathan et al. 2019) and CADD v1.6 (Kircher et al. 2014) (which incorporates SpliceAI predictions) were included in the assessment alongside the challenge models as a comparison to emerging industry standards. CADD-phred scores were obtained by uploading a VCF to the CADD webserver (https://cadd.gs.washington.edu/score). SpliceAI scores were obtained from Ensembl’s Variant Effect Predictor (VEP) web interface (McLaren et al. 2016) (44 variants scored) or using the SpliceAI webserver from the Broad Institute (https://spliceailookup.broadinstitute.org/, 11 variants that were not scored by VEP; options: hg38, masked scores, max distance 50bp). A cut-off of 0.2 was used for SpliceAI scores, and 18 for CADD.

## Results

### Variant characteristics of challenge set

Of the 56 variants in the final analysis, the majority (n=49, 87.5%) were SNVs, with 7 indels (12.5%). The variants fell within 42 different genes, broadly representative of clinical genetics referrals in the UK, with the majority of genes having a single variant in the set, and only 7 genes with >1 variant (*BRCA1* n=6, *FBN1* n=4, *MYBPC3* n=3, *BRCA2* n=2, *SCN5A* n=2, *APC* n=2, *USP7* n=2). 37 variants (66%) were found to affect splicing, while 19 (34%) had no observable impact.

Variants were divided into 5 groups by their positions relative to intron-exon boundaries. There were 16 variants within 10bp of a splice acceptor site (NearAcc), and 23 within 10bp of a splice donor site (NearDon). 10 exonic variants >10bp from either splice site were classed as Exonic>10. Intronic variants >10bp from their nearest splice site were termed Intronic Distant (six upstream of the acceptor, one downstream of the donor). The locations of all variants relative to the intron-exon boundary and whether the variants were determined to be splice disrupting or not are given in **Fig1**.

**Fig1.**
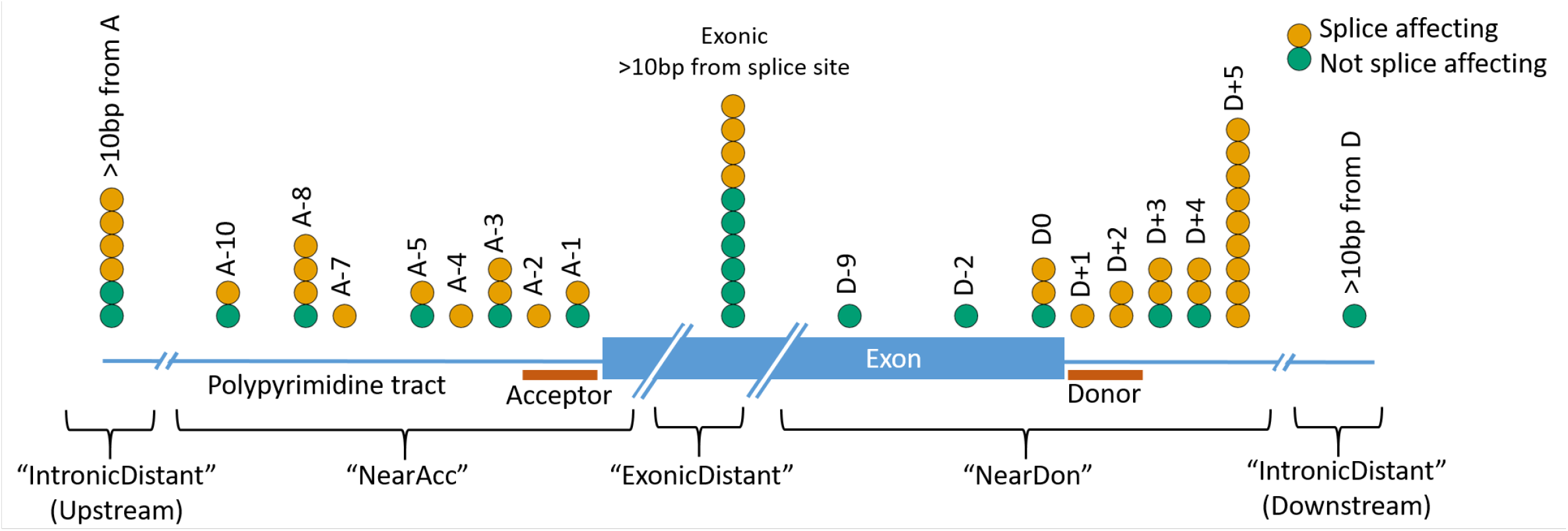
Schematic diagram showing locations of the 56 challenge variants in relation to their nearest splice site, with colour indicating whether (yellow) or not (green) each variant was determined experimentally to impact splicing.

### Challenge participants

Eight teams submitted predictions for the challenge, with two teams submitting predictions from multiple models, giving 12 models altogether. **Table 1** gives a summary of the approach taken by each model, which was provided by the challenge entrants upon submission of their predictions, but blinded to the assessors until after the assessment period.

**Table 1.**
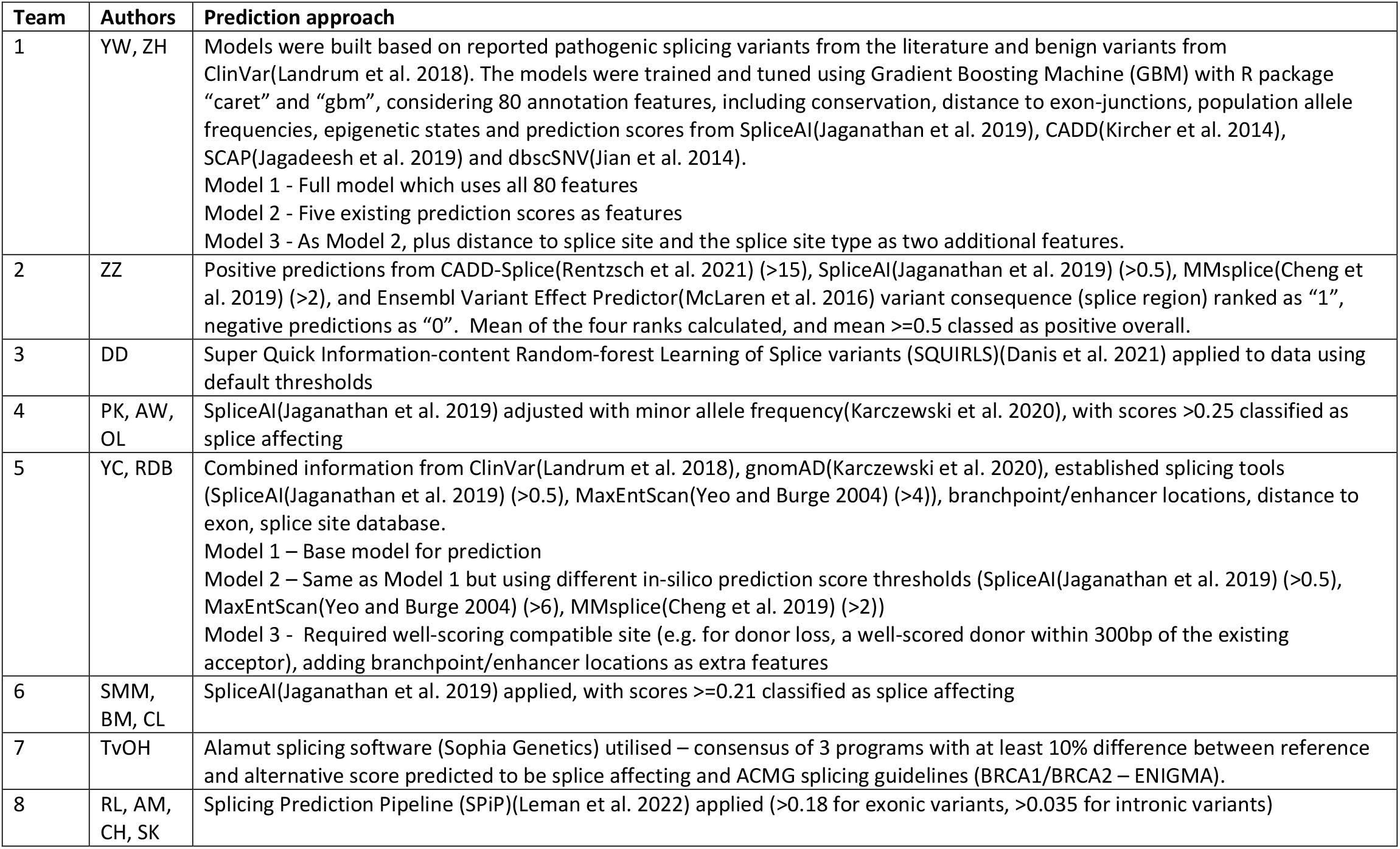
Summary of the prediction approaches of the 12 models from 8 entrants. Additional information on Teams 4 and 5 given in the **Supplementary Methods**.

### Model performance across 56 variants

**Table 2** summarises the performance metrics of the 12 models, along with CADD and SpliceAI. Full variant information, scores and binary predictions for the 12 models, SpliceAI and CADD and experimental outcome of splicing status are given in **Supplementary Table 1**. The ROC plots for each model are shown in **Fig2**, and **Supplementary Fig1** shows the performance of each method on each variant across the splicing region.

**Table 2.**
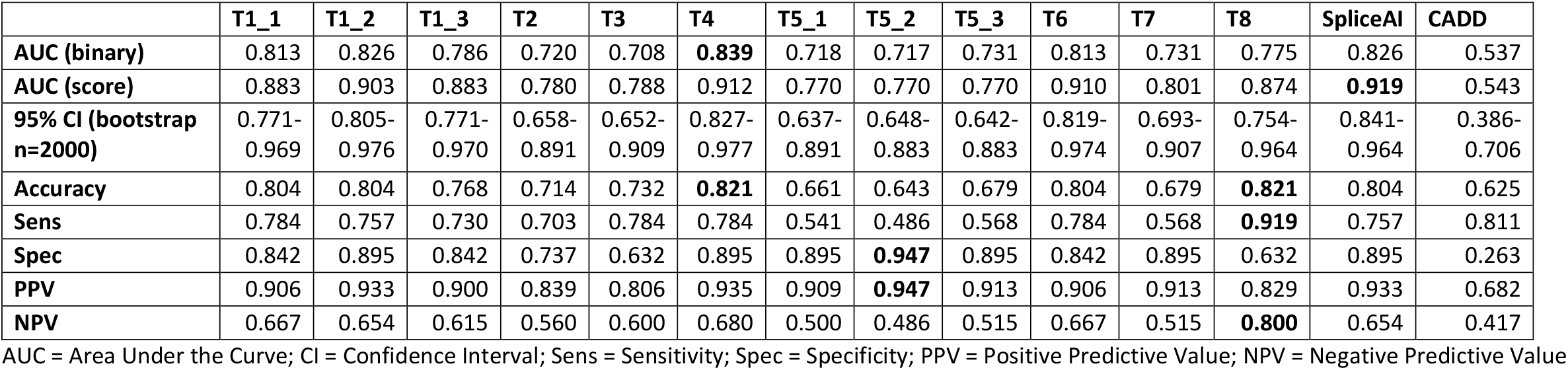
Summary statistics on predictive performance of the 12 competition entrants plus SpliceAI and CADD on the 56 challenge variants. Maximum value for each metric indicated in bold.

|  | T1_1 | T1_2 | T1_3 | T2 | T3 | T4 | T5_1 | T5_2 | T5_3 | T6 | T7 | T8 | SpliceAI | CADD |
| --- | --- | --- | --- | --- | --- | --- | --- | --- | --- | --- | --- | --- | --- | --- |
| **AUC (binary)** | 0.813 | 0.826 | 0.786 | 0.720 | 0.708 | **0.839** | 0.718 | 0.717 | 0.731 | 0.813 | 0.731 | 0.775 | 0.826 | 0.537 |
| **AUC (score)** | 0.883 | 0.903 | 0.883 | 0.780 | 0.788 | 0.912 | 0.770 | 0.770 | 0.770 | 0.910 | 0.801 | 0.874 | **0.919** | 0.543 |
| **95% CI (bootstrap n=2000)** | 0.771-0.969 | 0.805-0.976 | 0.771-0.970 | 0.658-0.891 | 0.652-0.909 | 0.827-0.977 | 0.637-0.891 | 0.648-0.883 | 0.642-0.883 | 0.819-0.974 | 0.693-0.907 | 0.754-0.964 | 0.841-0.964 | 0.386-0.706 |
| **Accuracy** | 0.804 | 0.804 | 0.768 | 0.714 | 0.732 | **0.821** | 0.661 | 0.643 | 0.679 | 0.804 | 0.679 | **0.821** | 0.804 | 0.625 |
| **Sens** | 0.784 | 0.757 | 0.730 | 0.703 | 0.784 | 0.784 | 0.541 | 0.486 | 0.568 | 0.784 | 0.568 | **0.919** | 0.757 | 0.811 |
| **Spec** | 0.842 | 0.895 | 0.842 | 0.737 | 0.632 | 0.895 | 0.895 | **0.947** | 0.895 | 0.842 | 0.895 | 0.632 | 0.895 | 0.263 |
| **PPV** | 0.906 | 0.933 | 0.900 | 0.839 | 0.806 | 0.935 | 0.909 | **0.947** | 0.913 | 0.906 | 0.913 | 0.829 | 0.933 | 0.682 |
| **NPV** | 0.667 | 0.654 | 0.615 | 0.560 | 0.600 | 0.680 | 0.500 | 0.486 | 0.515 | 0.667 | 0.515 | **0.800** | 0.654 | 0.417 |

177 AUC = Area Under the Curve; CI = Confidence Interval; Sens = Sensitivity; Spec = Specificity; PPV = Positive Predictive Value; NPV = Negative Predictive Value

**Fig2.**
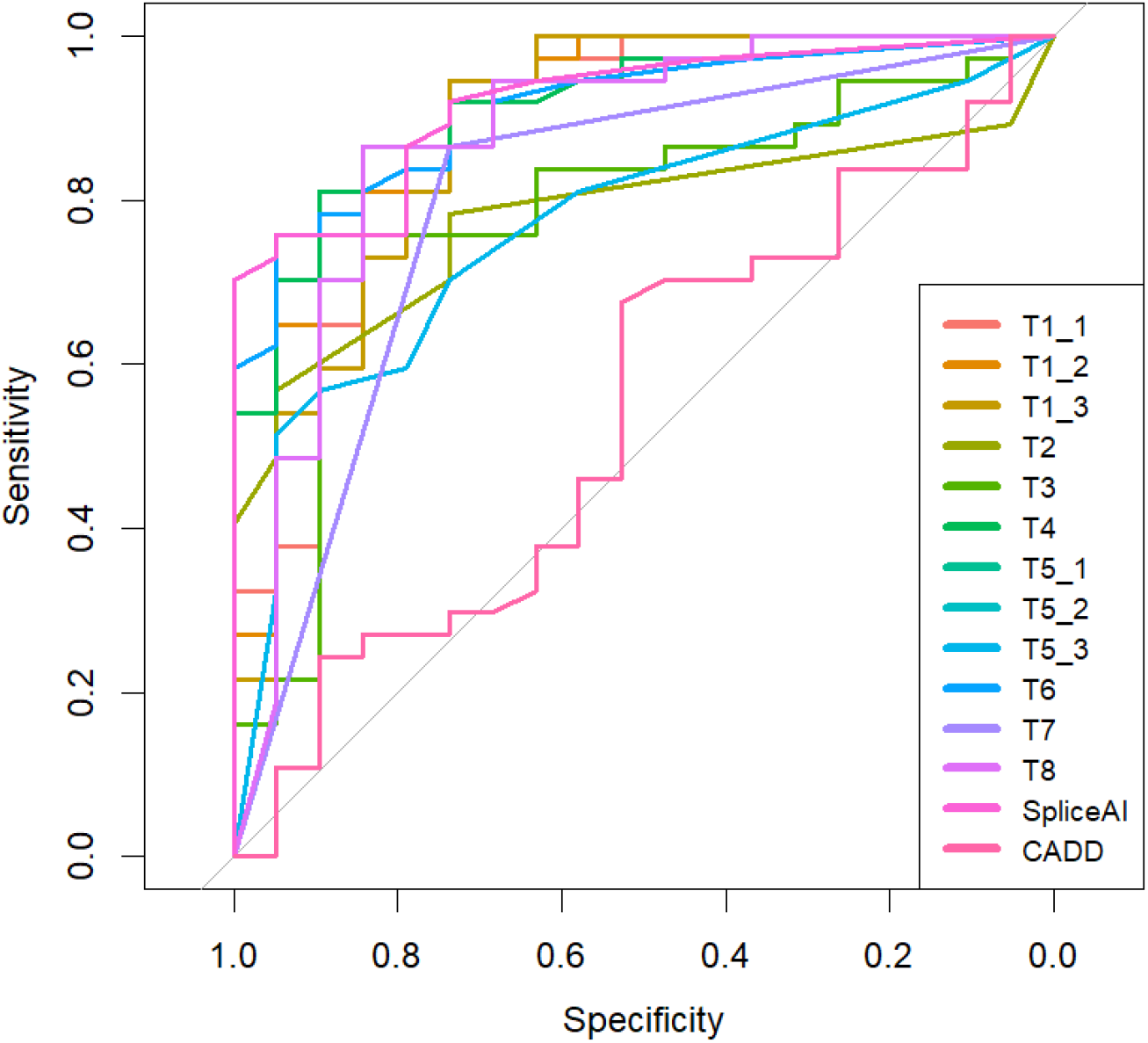
Receiver operating characteristic (ROC) curves of model performance based on prediction scores. For Area Under Curve (AUC), see **Table 2**.

No single approach performed optimally on all assessment metrics (**Table 2**). Overall accuracy was joint highest in Teams 4 and 8 at 0.82, with Team 4 also achieving the highest binary outcome AUC at 0.839 (**Fig2**). Team 8 ranked highest on the related metrics for sensitivity (0.919) and NPV (0.800), indicating its permissive prediction approach. Conversely, Team 5’s Model 2 performed the best in terms of specificity (0.947) and PPV (0.947), with the lowest proportion of false positive findings. All three models by Team 1, plus Team 4 and Team 6 achieved over 70% in both sensitivity and specificity, indicating more balanced performance.

Included as comparators were SpliceAI with a cut-off of 0.2 and CADD with a cut-off of 18. SpliceAI was competitive with the challenge entrants, ranking near-top but not top on all metrics, and indeed top in the AUC when measured using prediction score rather than binary prediction outcome. CADD, however, performed poorly on the challenge set with specificity in particular being very low (0.263).

### Performance comparison by variant type

In order to get an overall impression of the performance of the methods on different types of variants, variants were grouped by location relative to their nearest splice site (**Fig3**), as described in Methods. All methods performed better on exonic distant variants than intronic distant variants, with the exception of SpliceAI, which correctly predicted all seven intronic distant variants. Across methods, there was a high degree of consistency in the proportion of variants correctly predicted in the near acceptor region, and a high degree of variance in performance in the intronic distant set. The types of error differed across regions, with the near acceptor region and exonic distant region having very few false positive predictions across all methods, while almost all methods gave false positive predictions in the near donor and intronic distant regions (**Supplementary Fig2**).

**Fig3.**
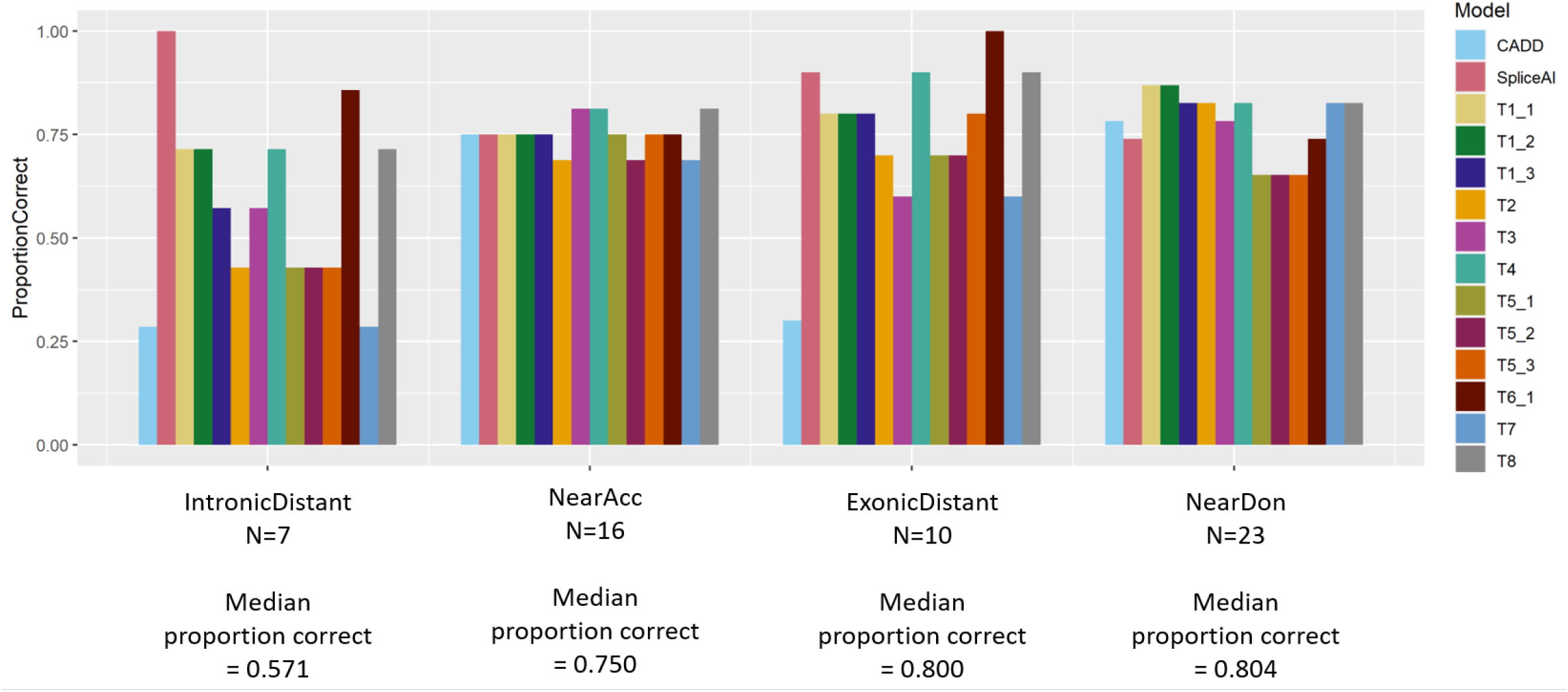
Proportion of variants correctly predicted by each method in the different regions (near acceptor, near donor, exonic and intronic distant).

**Fig4.**
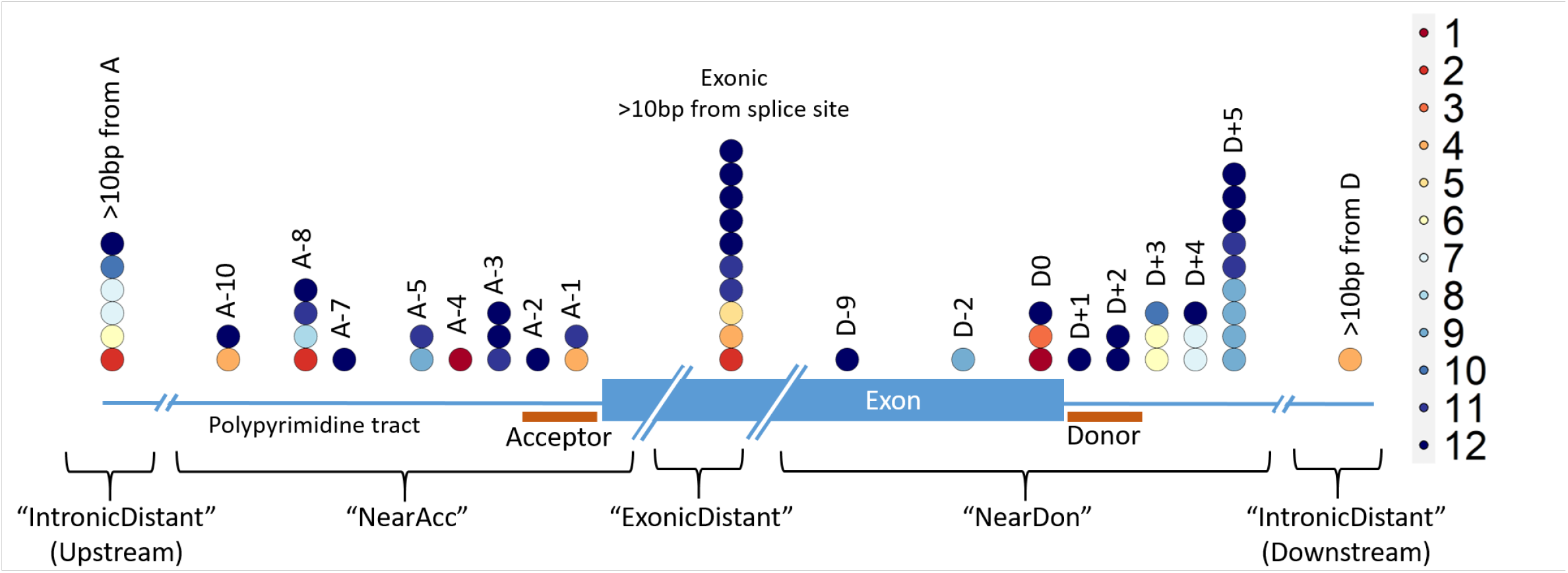
Variants across the splicing region coloured by the number of prediction methods (out of the 12 challenge entrants) that correctly predicted the splicing outcome.

We also compared the performance of the approaches on SNVs vs indels, and found all methods except CADD had higher accuracy on SNVs than indels (**Supplementary Fig3**).

### Some variants are consistently mispredicted

21 of the variants (37.5%) were correctly predicted by all 12 submitted prediction methods. None of the variants were incorrectly predicted by all methods, but 14 variants (25%) were predicted correctly by <=50% of the methods, with two variants only being correctly predicted by a single method. These were a splice-affecting single nucleotide deletion 4bp from a splice acceptor site in *KANSL1* (correctly predicted by Team 3) and an SNV in the last base of an exon in *TRPM6* which despite altering the conserved last G nucleotide did not affect splicing in functional testing (correctly predicted by Team 4).

## Discussion

The CAGI6 Splicing VUS challenge assessed the performance of 14 prediction approaches on a set of 56 clinically relevant variants whose impact on splicing had been functionally tested using RT-PCR. A variety of approaches were adopted, and several methods equalled or exceeded the performance of the emergent field leader, SpliceAI.

While Teams 4 and 8 had joint highest overall accuracy, there was no single optimal method for the Splicing VUS challenge, since several different models performed optimally on different metrics. Choice of approach may therefore be dependent on the specific nature of the predictions required. Seeking a molecular diagnosis for a particular family may favour sensitivity over specificity, since overlooking a causal variant would prevent this aim, so Team 8’s approach with almost 92% sensitivity may be preferred. Seeking confident splice disrupting candidates for functional validation or mechanistic research may call for greater specificity than sensitivity to avoid wasting resources on false positive variants that do not have an impact, in which case Team 5’s model 2 with almost 95% specificity may be the strategy of choice.

SpliceAI and CADDv1.6 were chosen as comparators for the entrants to the Splicing VUS challenge and were run by the assessors on the 56 challenge variants. SpliceAI has been emerging as a field leader in recent years, with accuracies >90% attained in several studies (Wai et al. 2020; Ha et al. 2021; Strauch et al. 2022), although variable performance reported by some (Riepe 2020) which is more consistent with our observed 80.4% overall accuracy in this study.

CADD did not perform well on the challenge variants, achieving an overall accuracy of 62.5%. However, this was predominantly driven by a very low specificity, which is to be expected from CADD, since it is not only the impact on splicing being assessed by the tool, but overall deleteriousness. For example, missense variants which were not found to affect splicing in the challenge set may still have been pathogenic through impact on protein structure and/or function. For such variants, CADD would accurately classify them as deleterious in general, but in our assessment solely of splicing impact, this would appear as a false positive, lowering CADD’s specificity. Notably, the version of CADD included in the assessment (v1.6) includes SpliceAI and additional splicing prediction tools in its underlying model (Rentzsch et al. 2021). Scoring the challenge variants with CADD v1.5 which did not include these splicing metrics resulted in an overall accuracy around 44.6% (data not shown). From these values it is clear that the explicit inclusion of splicing prediction methods within CADD’s underlying model has improved its ability to predict variants that impact splicing. CADD’s broad approach makes it a versatile tool for prediction of deleteriousness for many different variant types. At present, however, if predicting a variant’s splicing impact is the sole aim, the use of more specialised splicing tools is more appropriate.

Of note, SpliceAI featured heavily across the predictive strategies, being the sole predictive method for Team 6 and contributing heavily to the predictions of Team 4, which were weighted by MAF, as well as being run as a comparator by the assessors. Differences in the performance of these approaches highlight that even with the same nominal method, there can be variance in predictions depending on how the scores are obtained, and the thresholds that are used to determine positive predictions. There were just three approaches that did not include SpliceAI as part of their predictions, two utilising instead recent machine learning based prediction tools SQUIRLS (Danis et al. 2021) and SPiP (Leman et al. 2022), and one based on the splicing prediction tools available within the Alamut software, which has been widely used in clinical practice. Of the three, SPiP was the only method to achieve greater accuracy than SpliceAI.

A major strength of the challenge in terms of providing a real-world assessment of the performance of these tools is the ascertainment of the challenge variants from genuine clinical practice, where potential splice altering variants in genes relevant to the patient’s presentation were referred for validation. This is precisely the type of variant splicing prediction models should be tested on when assessing their suitability for clinical application in rare disorders. It highlights that even in exceptionally well-studied genes, such as the BRCA genes, challenges in variant interpretation remain, since 3 of 8 variants across *BRCA1* and *BRCA2* were incorrectly predicted by over half of challenge methods, and only two of these were accurately predicted by all methods. However, the relatively small sample size makes it difficult to draw any major inferences and is a significant limitation of the study. Apparent variance in performance may be stochastic at such a sample size, and may not be fully reflective of overall performance in a wider context. It also made drawing firm conclusions about performance in subsets of the data, e.g. split by location, variant type, or disease group challenging. However, ascertaining a large body of clinical variants, validating the splicing impact and keeping that private, as is needed for a blinded challenge such as the CAGI6 Splicing VUS challenge, raises ethical concerns. Accurate and timely variant interpretation is reliant on sharing of data, and withholding a large body of functionally validated variants from resources such as ClinVar (Landrum et al. 2018) which are heavily used in clinical assessment of variants does not represent good practice.

This small but highly clinically relevant challenge assessed the performance of 12 prediction methods plus SpliceAI and CADD on 56 clinically ascertained variants and found SpliceAI weighted by allele frequency and SPiP to be the most accurate overall, while other methods had particular strengths in their sensitivity or specificity. A quarter of variants were incorrectly predicted by half or more of the methods, showing there is still improvement to be made. Furthermore, this challenge was limited to a binary outcome – whether or not splicing was disrupted, but did not address the nature of that disruption, which may present an even greater challenge. A larger assessment set that would enable further investigation of the types of variants that are consistently incorrectly predicted may help direct efforts for refinement of models moving forwards.

## Acknowledgements

We thank the CAGI organisers for their commitment to improving variant interpretation and for making this challenge happen. The CAGI experiment is supported by NIH U24 HG007346.

We acknowledge the NIHR Clinical Research Network (CRN) in recruiting the participants and the Musketeers Memorandum, as well as support from the NIHR UK Rare Genetic Disease Consortium. The authors acknowledge the use of the IRIDIS High Performance Computing Facility, and associated support services at the University of Southampton, in the completion of this work.

For the purpose of open access, the author has applied a CC BY public copyright licence to any Author Accepted Manuscript version arising from this submission.

## Funding

The Baralle Lab is supported by the NIHR Research Professorship awarded to D.B. (RP-2016–07-011). JL is supported by an Anniversary Fellowship from the University of Southampton. Some of the functional validations of variants were funded by a Wessex Medical Research Innovation Grant awarded to JL. RDB is supported by a New South Wales Health Cardiovascular Disease Senior Scientist Grant.

## Author contributions

DB and JL conceived of the challenge. AGLD, DJB and JL selected variants to include in the set, which had been functionally validated by HAW and DJB. JL assessed challenge entrants and conducted data analysis. CJO conducted additional analyses and presented the findings at the CAGI6 conference. All further authors submitted prediction methods in response to the challenge. JL drafted the manuscript, with revision suggestions and final approval from all other authors.

## Data availability

All data generated or analysed during this study are included in this published article [and its supplementary information files].

## Notes

### Competing Interest Statement

The authors have declared no competing interest.

