## Supplementary Information and Figures for "Predicting the impact of rare variants on RNA splicing in CAGI6"

**Supplementary Methods**

**Team 4**

The submission was based on SpliceAI scores and the minor allele frequency (MAF) of the variants.

However, in practice the outcome mostly reflected the SpliceAI scores.

Specifically:

We used the precomputed SpliceAI scores. These scores consisted of four values (dAG, dAL, dDG, dDL; d=delta, A=acceptor, D=donor, G=gain, L=loss).

We combined these values as: overall SpliceAI score = 1-(1-dAG)(1-dAL)(1-dDG)(1-dDL)

This overall SpliceAI score was combined with a weighting factor derived from the MAF of variants according to the gnomAD Genomes v3.1.1,  gnomAD Exomes r2.1.1, and UK Biobank (200k) databases.

This weighting factor considered a minimum MAF (half of the MAF that corresponds to singletons) and was equal to the log(MAF_variant)/log(MAF_min).

The final score was computed as the product of the overall SpliceAI score with the smallest weighting factor corresponding to any of the three MAF databases.

Finally, we used a threshold of 0.25 to separate pathogenic from benign effects.

**Team 5**

**Brief description of the models**

Each variant is annotated for the genomic location and consequence using Varcode, v0.5.15, and the gene's inheritance mode (autosomal or X-linked dominant or recessive) as retrieved from the OMIM database. Variants are flagged as rare if found <15 times in the gnomAD database for dominant genes and <1500 times for recessive genes. A variable indicates whether a variant is reported as pathogenic or likely pathogenic in ClinVar database. *In silico* splice prediction scores are added as separate parameters from three tools (MaxEntScan [1], SpliceAI [2], and MMSplice [3]) with specific cut-off values of 4 for the absolute difference between reference and wildtype alleles for MaxEntScan and >=0.5 for SpliceAI and MMSplice predictions.

In our second model, additional parameters are incorporated to enhance prediction of variants that create new splice donor or acceptor sites. These include whether there is an existing and corresponding donor or acceptor site within 300bp of the newly created donor or acceptor site. For example, if the variant creates a splice donor site, we search the upstream 300bp for any acceptor sites with a MaxEntScan score >4. We also add a variable for whether there is an exon enhancer sequence [GAAGA; TGCGTC; CACACGA; GGCCCCTG; TCACAGG] within the pseudo exon [4], and whether there is a branch point motif within 50bp upstream of an acceptor site.

To train the model, we randomly selected 100 pathogenic splice site variants and 200 missense variants classified as having uncertain clinical significance from the ClinVar database. We fit a random forest regression [5] to predict the variants that alter splicing. After fitting the model, we generate a result score, ranging from 0 (less likely) to 1 (extremely likely), indicating whether the variant disrupts splicing.

Our models are limited by the relatively small size of the training data set and the dependence on the robustness of ClinVar classifications. Most splice site variants were in canonical splices. Nonetheless, the inclusion of parameters such as corresponding sites and multiple prediction tools make our approach comprehensive and useful for predicting splice-disruptive variants.

**Supplementary Figures**

**Supplementary Fig1.** Experimentally validated splicing outcome and prediction accuracy for all 56 variants across the splicing region.

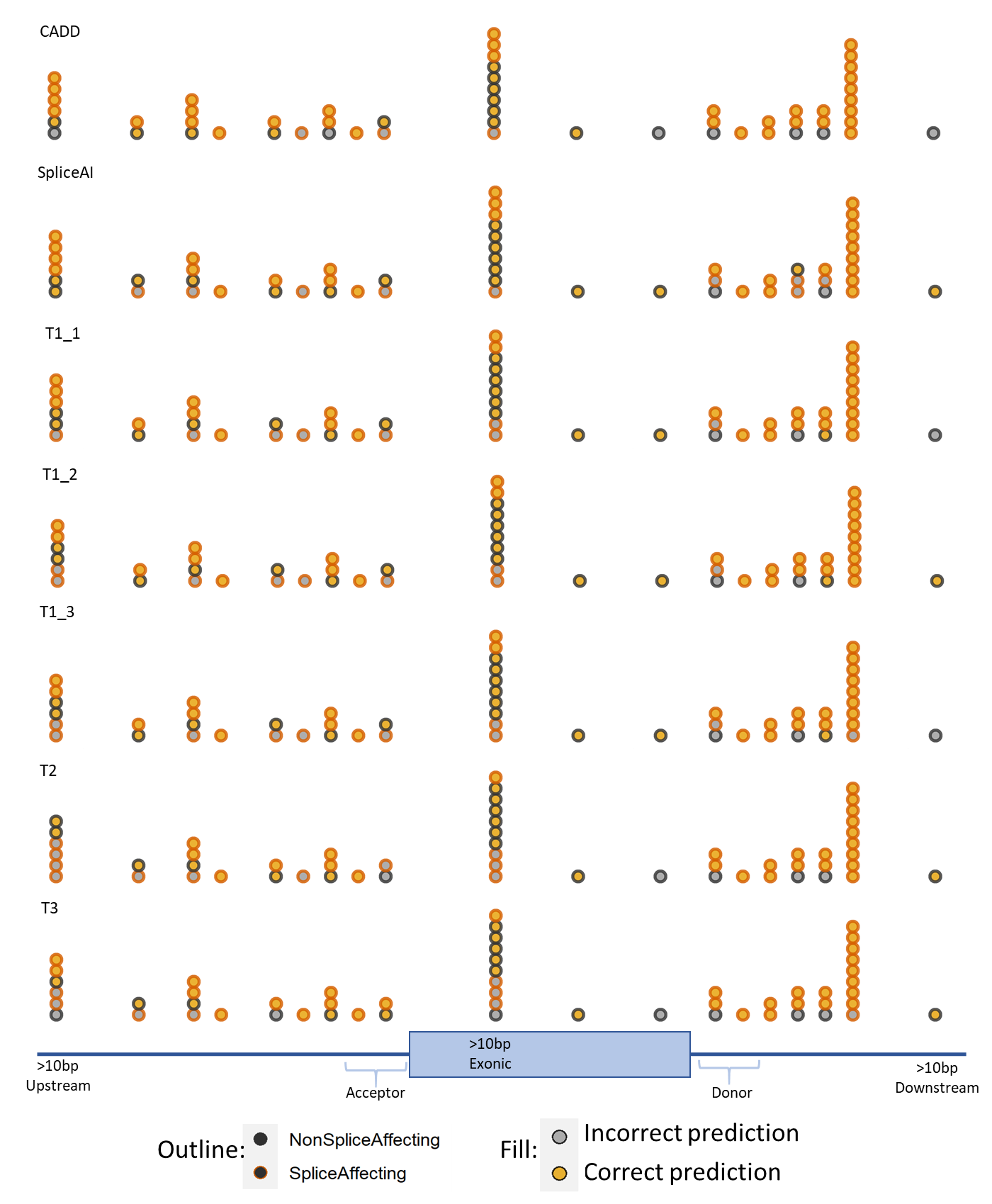

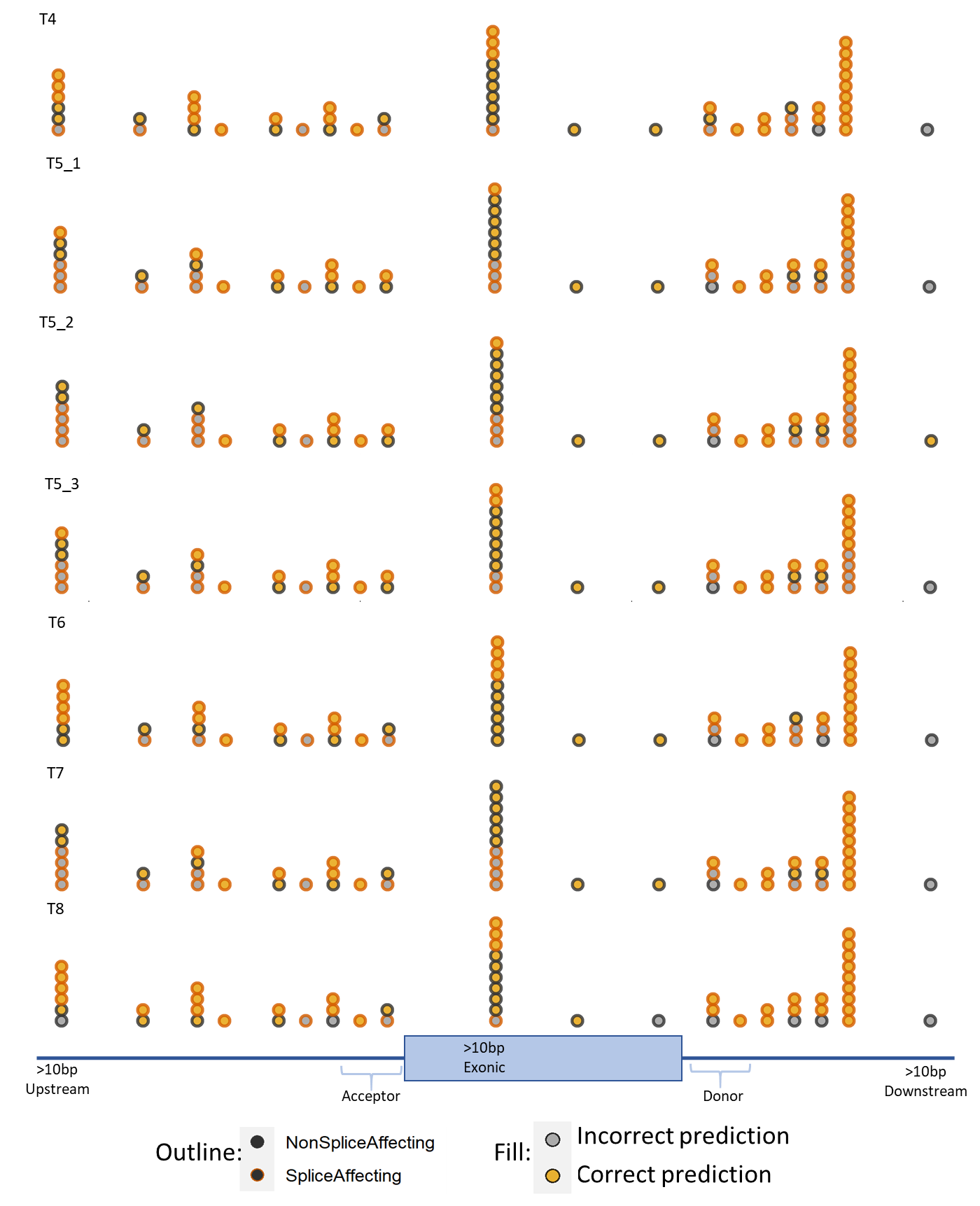

**Supplementary Fig2.** Breakdown of false negative (FN), false positive (FP), true negative (TN) and true positive (TP) predictions for the 12 challenge entrants plus CADD and SpliceAI grouped by region (near acceptor, near donor, exonic and intronic distant).
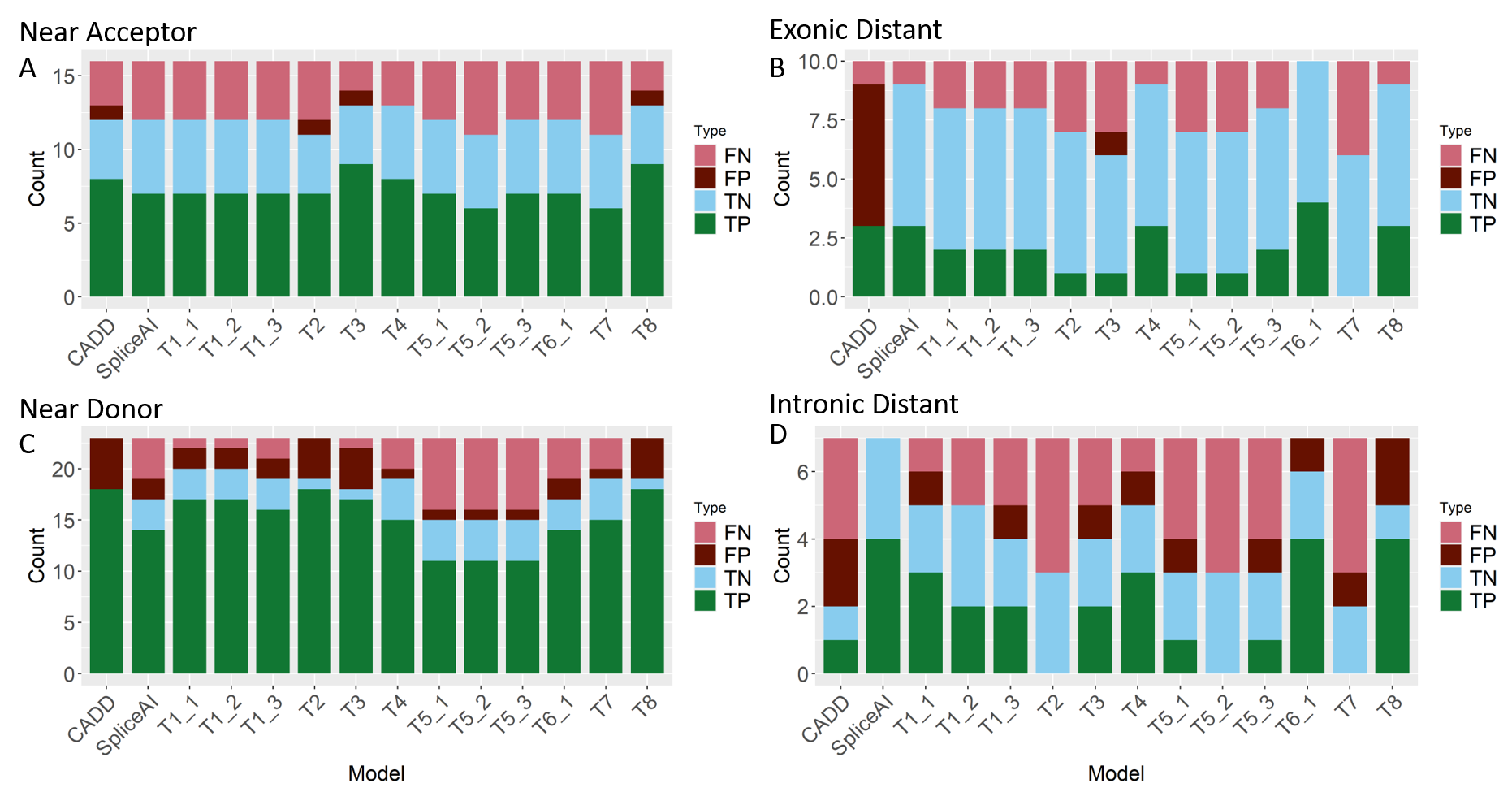

**Supplementary Fig3.** Proportion of correctly predicted splicing outcomes for each of the 12 models plus SpliceAI and CADD split between SNVs and indels.

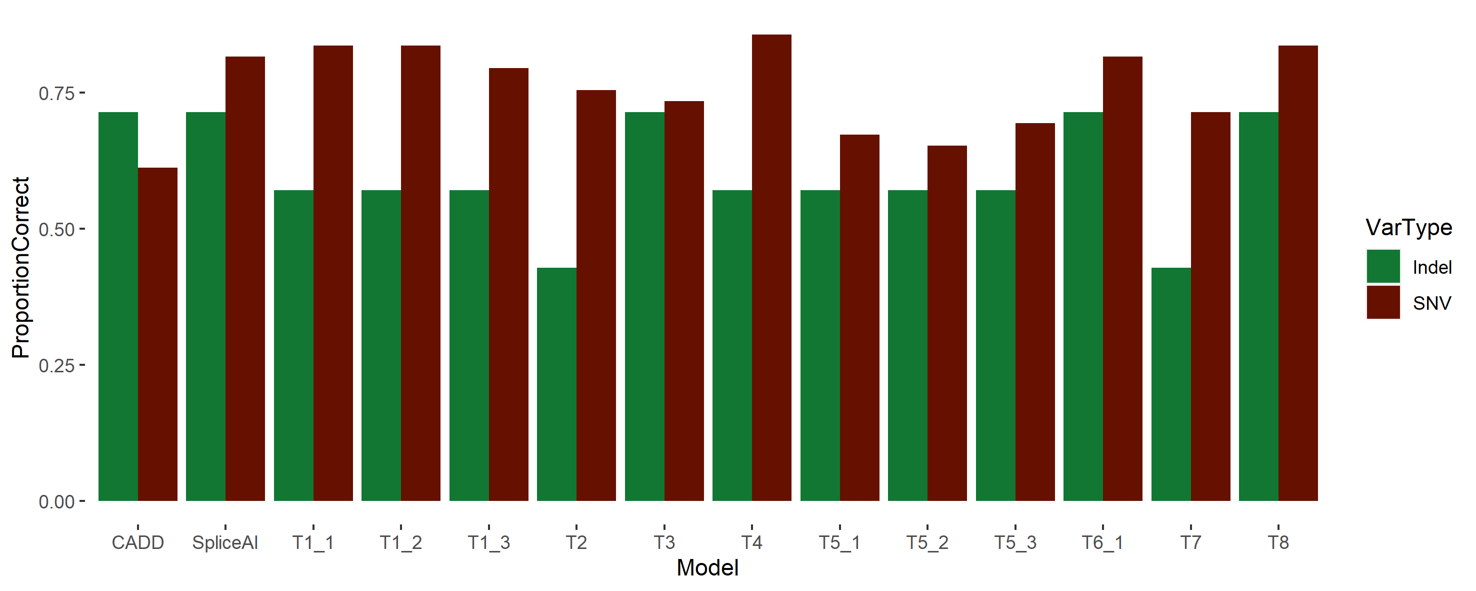
